# Internally coupled middle ears enhance the range of interaural time differences heard by the chicken

**DOI:** 10.1101/512715

**Authors:** Christine Köppl

**Affiliations:** Department of Neuroscience, School of Medicine and Health Sciences, Carl von Ossietzky University Oldenburg, 26129 Oldenburg, Germany; Cluster of Excellence “Hearing4all” and Research Center Neurosensory Science, Carl von Ossietzky University Oldenburg, 26129 Oldenburg, Germany

**Keywords:** auditory, hearing, sound localization, bird, avian, ITD

## Abstract

Interaural time differences (ITD) are one of several principle cues for localizing sounds. However, ITD are in the sub-millisecond range for most animals. Because the neural processing of such small ITDs pushes the limit of temporal resolution, the precise ITD-range for a given species and its usefulness - relative to other localization cues - was a powerful selective force in the evolution of the neural circuits involved. Birds and other non-mammals have internally coupled middle ears working as pressure-difference receivers that may significantly enhance ITD, depending on the precise properties of the interaural connection. Here, the extent of this internal coupling was investigated in chickens, specifically under the same experimental conditions as typically used in neurophysiology of ITD-coding circuits, i.e. with headphone stimulation. Cochlear microphonics (CM) were recorded simultaneously from both ears of anesthetized chickens under monaural and binaural stimulation, using pure tones from 0.1 to 3 kHz. Interaural transmission peaked at 1.5 kHz at a loss of only −5.5 dB; the mean interaural delay was 264 μs. CM amplitude strongly modulated as a function of ITD, confirming significant interaural coupling. The “ITD heard” derived from the CM phases in both ears showed enhancement, compared to the acoustic stimuli, by a factor of up to 1.8. However, the closed sound delivery systems impaired interaural transmission at low frequencies (< 1 kHz). We identify factors that need to be considered when interpreting neurophysiological data obtained under these conditions, and relating them to the natural free-field condition.

**Summary statement:** The interaural time differences that chickens can use for sound localization are significantly greater than their small head size suggests. Closed-system sound stimulation can, however, produce complex artefacts.

## Introduction

Localization of sounds originating in the environment is performed without effort by humans and many animals. This apparent ease belies the complexity of the underlying physical and neurophysiological processes. There is a number of principle cues - interaural time and level differences in azimuth and spectral composition in elevation – but their availability and relative usefulness are highly dependent on the size of the animal and its frequency range of hearing (e.g., Köppl, 2009). In the low-frequency range, typically up to a few kHz, interaural time differences (ITD) are the best cue to azimuth (e.g., Hartmann, 1999). However, for all but the largest animals, ITDs remain below 1 ms and thus represent a challenge for the nervous system to encode timing and determine the interaural difference with appropriate precision. Although it is undisputed that humans and other animals with good low-frequency hearing rely on ITD for sound localization in azimuth (e.g., (Brown and May, 2005), the neural mechanisms underlying this are less clear. Several mechanisms of encoding ITD have been suggested, with good experimental evidence for each, in different species, and sometimes even in the same species (reviews in Ashida and Carr, 2011; Grothe et al., 2010; Joris and Yin, 2007; Vonderschen and Wagner, 2014). This naturally raises the question as to the constraints and specific conditions that might have favored the evolution of different mechanisms (Carr and Christensen-Dalsgaard, 2015; Carr and Christensen-Dalsgaard, 2016; Grothe and Pecka, 2014; Köppl, 2009). The precise range of ITDs available to an animal is an important argument in this discussion, but wrong assumptions have often been made about this.

The acoustic ITD appears to be straightforward to predict if the size of the head is known and the head is approximated as a sphere (Kuhn, 1977). At low frequencies, the maximal ITD arising from a sound source 90° to one side is 3r/v (where r is the radius of the sphere and v the speed of sound). However, actual measurements in a range of animal species have since shown that the acoustic ITD between the outsides of both eardrums is always larger than this prediction, typically by a factor of about 1.5 (cat: Tollin and Koka, 2009; guinea pig: Sterbing et al., 2003; gerbil: Maki and Furukawa, 2005; chinchilla: Jones et al., 2011; barn owl: Hausmann et al., 2010; Poganiatz et al., 2001; von Campenhausen and Wagner, 2006). Recently, this was also confirmed for the chicken (Schnyder et al., 2014; estimated from phase measurements shown in their Supplemental Fig. 9). Thus classic assumptions about the ITD range experienced by an animal and based on a spherical head model, need to be revised upwards.

There is more to this issue. Both mammalian and avian species are prominent animal models for investigating the neural processing mechanisms of ITD. However, little attention has been paid in this context to a salient difference in their middle ears that has a potentially crucial impact on ITD processing. Unlike mammalian ears, the middle ears of birds are acoustically connected through skull spaces, often collectively termed the interaural canal. This internal coupling turns the ears into pressure-difference receivers, with sound reaching each eardrum from both sides. The driving force is then the instantaneous pressure difference across the eardrum, and the phase of eardrum movement is the difference between the phases of the direct and indirect component, weighed by the interaural transmission gain. Importantly, depending on the physical dimensions of the head, the sound wavelength, and the attenuation across the interaural connections, increased directional cues to sound location may be generated, including enhanced ITDs (Christensen-Dalsgaard, 2011; Michelsen and Larsen, 2008).

The presence of internal connections between the middle ears of birds (and, more generally, archosaurs) was demonstrated early and is undisputed (e.g., Owen, 1850; Schwartzkopff, 1952; Wada, 1924). However, the presence of internal coupling between the middle ears is merely a prerequisite and in itself does not prove that significant directional cues arise from it. It is the precise degree of interaural transmission that determines whether a significant directionality actually results. These details have proven difficult to define. The morphology of the connections across the head remains ill-characterized, in large part due to the extensively pneumatized and trabeculated structure of avian bones, which generates a myriad of potential skull paths. Connections between the two sides likely include more than the classic ventral “interaural canal” (Bierman et al., 2014; Christensen-Dalsgaard, 2011; Larsen et al., 2016; Rosowski, 1979). Attempts to quantify the physiological effect of internal coupling in birds include acoustic measurements at various locations both outside and inside the skull (Hill et al., 1980; Rosowski, 1979; Rosowski and Saunders, 1980), and measurements of eardrum vibration or recordings of cochlear microphonics as a proxy for eardrum vibration (Calford and Piddington, 1988; Hyson et al., 1994; Klump and Larsen, 1992; Larsen et al., 2006; Lewald, 1990; Moiseff, 1989; Rosowski, 1979). Conclusions about the significance of interaural connections varied widely (reviewed by Christensen-Dalsgaard, 2005; Klump, 2000), no doubt further complicated by the discovery of a major source of experimental artefact, the buildup of negative middle-ear pressure under anesthesia (Larsen et al., 2016; Larsen et al., 1997).

The present study aimed to re-investigate the effect of internally coupled ears in the chicken, with a specific emphasis on ITD. The chicken is a well-studied animal model in the context of neural ITD coding. The possibility of internal coupling of the ears raises a serious problem for the controlled presentation of ITD, which is typically done via headphones when testing neural selectivity for ITD: In this situation, the acoustically presented ITD may not be the ITD heard by the bird, and this confounds the interpretation of neural responses. It is therefore important to quantify the effect of internal coupling of the ears for the species in question.

## Material and Methods

### Animal anesthesia and homeostasis

Cochlear microphonics were recorded in 8 chickens (*Gallus gallus domesticus*) of commercial egglayer breeds, aged posthatching day (P) 28 to 37, and weighing between 100 and 200g. Their head widths, measured with calipers between the entrances to the ear canals, were 22-23 mm. Chickens were deprived of food for at least 2 hours, in preparation for anesthesia that was initiated by injecting 20mg/kg ketamine hydrochloride and 3mg/kg xylazine intramuscularly. Supplementary doses were adjusted individually, at 50 – 100 % of the initial, usually every 30 – 50 minutes. The primary monitor for depth of anesthesia was a combined EKG- and muscle-potential recording via insect needles inserted into the muscles of a leg and the contralateral wing. This signal was amplified (Grass P15) and constantly displayed on an oscilloscope. Cloacal temperature was held constant at 41.5° via a feedback-controlled heating blanket (World Precision Instruments, Sarasota, USA) wrapped around the chicken’s body. The trachea was exposed in the neck region, cut and intubated with a short piece of matching tubing to prevent problems from salivation; through this, chickens breathed normal room air unaided. The chicken’s head was wrapped with strips of plaster-of-Paris, which was connected to a metal head holder by dental cement, to fix the head in a defined position.

### Electrode placement and recordings

Bilateral surgical openings through the neck muscles and underlying bone provided access to the middle-ear spaces. Electrodes custom-made of insulated silver wire with a small bare silver ball at the end were inserted and the silver ball placed onto the membrane covering the recessus scalae tympani. In a few cases, the membrane was slit and electrodes inserted into scala tympani. This increased the recorded CM amplitudes somewhat but did not provide sufficient advantage to adopt routinely. Electrodes were glued into place on the skull’s surface with tissue glue and dental cement. Reference electrodes were placed under the skin nearby and were either silver ball electrodes of the same type, separate for left and right (4 experiments) or an Ag/AgCl pellet shared for both channels (4 experiments). The surgical holes were left open during all measurements, thus ensuring middle-ear ventilation. Signals were amplified ×500,000 by a Tucker-Davis Technologies (TDT, Alachua, USA) DB4 amplifier, bandpass filtered at 100 Hz to 15 kHz, and the two channels fed to the inputs of a TDT DD1 A/D converter that was connected to a TDT AP2 signal processing board. Data acquisition of the analog waveforms was controlled by custom-written software (“XDPHYS” by the laboratory of M. Konishi, Caltech, USA).

### Sound stimulation

Sound stimulation was through custom-made closed sound systems placed at the entrance of both ear canals. They contained a standard earphone (Sony MDR-E818LP) and calibrated miniature microphone (Knowles EM 3068) each. Microphone signals were amplified 40 dB by a custom-built amplifier. Sound-pressure levels and phases were calibrated individually at the start of each experiment and the calibrations used to adjust stimulus presentation online by custom-written software (xdphys, Caltech). Near-constant sound pressures down to the lowest frequency of 100 Hz suggested closed-system conditions, although no sealing agents were applied. Sealing was likely achieved through the feathers surrounding the ear canals. Stimuli were generated separately for the two ears using a TDT AP2 signal processing board. Both channels were fed to the earphones via D/A converters (TDT DD1), anti-aliasing filters (TDT FT6-2) and attenuators (TDT PA4). Stimuli were tone bursts of 50ms duration (including 5ms linear ramps), presented at a rate of 5/sec.

### Data collection and analysis

Monaural stimulation was usually tested at 8 standard frequencies (100, 333, 571, 1000, 1515, 2000, 2500 and 3030 Hz), at 40 to 80 dB SPL, in 10 dB steps. Responses to 200 repetitions of each stimulus were recorded.

The same standard frequencies were also tested binaurally, usually at two levels, 50 and 70 dB SPL. With binaural stimulation, ITD was also varied, within ± one stimulus period, in 10 steps per period. Repetitions were reduced from 200 to 50 for the higher level.

Recordings of the analog waveforms from left and right ears were always obtained simultaneously, regardless of whether the stimulation was monaural or binaural. An averaged analog response waveform was derived for each stimulus condition and contained both the compound action potential (CAP) and the cochlear microphonic (CM). Only the steady-state response between 15 and 45 ms re. stimulus onset was used for analysis, thus minimizing the neural component. A cosine function at the stimulus frequency was fitted and the amplitude and phase of this fit taken as the CM amplitude and phase. To eliminate recordings of insufficient signal-to-noise ratio, the fit amplitude was divided by the standard deviation of the averaged waveform *√2. The value of the resulting index is 1 if the waveform is identical to the fitted cosine and becomes zero if the waveform contains no stimulus frequency component (Köppl and Carr, 2008). Data were discarded if this index was below 0.5 for monaural recordings or, for binaural recordings, if it remained below 0.5 at all ITDs tested. If the CM amplitudes in binaural recordings showed an appreciable modulation with ITD, this ITD function was then fitted with a cosine function at the respective stimulus frequency (Viete et al., 1997) to determine peak ITD, defined as the peak closest to zero ITD.

### Acoustic measurements

In 3 chickens, the readings of the microphones integral to our sound systems were recorded under selected stimulation conditions by feeding their output (instead of the electrode recordings) into the A/D converter. Data analysis was exactly analogous to the procedures described above for CM recordings. The noise level of the microphones, estimated as the SPL where the readings exceeded the S/N criterion of 0.5 (see previous section) was 30 – 35 dB SPL.

### Blockage of interaural connections

In the same 3 chickens, an attempt was also made to block interaural connections. The ear canal on one side was widened through a small skin cut to gain access to the eardrum. The eardrum was pierced with a syringe loaded with petroleum jelly and jelly injected slowly behind the eardrum. The jelly appeared to liquefy quickly at the birds’ normal body temperature. Injection was stopped when the jelly began to exude to the outside of the eardrum. The sound system was re-positioned, both sides were re-calibrated and selected measurements repeated. In 2 of the 3 chickens, the skin cut was closed again with tissue glue in order to restore the ear canal as much as possible. Since these manipulations potentially not only blocked the interaural connections but also damaged the middle and inner ear on the manipulated side, only recordings of the unmanipulated ear were subsequently used. After euthanasia at the conclusion of the experiments, the chickens’ heads were placed in a refrigerator overnight to solidify the petroleum jelly. Care was taken to keep the head’s spatial orientation unchanged. Placement of the petroleum jelly was visualized by dissection on the next day.

## Results

### Dependence of monaural CM measurements on sound level

CM recordings under monaural stimulation were obtained at several sound levels, generally between 40 and 80 dB SPL, in 10 dB steps. In the ipsilateral ear, CM amplitudes were mostly above our criterion for S/N ratio at all those levels, i.e. the thresholds were 40 dB SPL or lower. Ipsilateral CM amplitude increased in a nearly linear fashion between 50 and 80 dB SPL at all frequencies (Fig. 1, top row of panels). In order to remain within this dynamic range in which CM amplitude was thus a reliable indicator of relative sound level, all comparisons between ipsi- and contralateral CM readings reported below were made at 70 dB SPL stimulus level. CM amplitudes in the two ears of a given animal, and at a given sound level, were generally similar. However, if there was an asymmetry, for unknown reasons there was an overall bias for higher amplitudes in the left ear. Comparisons between ipsi- and contralateral CM readings were therefore consistently carried out between matched recordings of the same ear to stimulation from the ipsi- and contralateral side, respectively (as opposed to simultaneous readings of the two CM recorded to stimulation of a given ear; schematically illustrated in Fig. 2A).

**Fig. 1:**
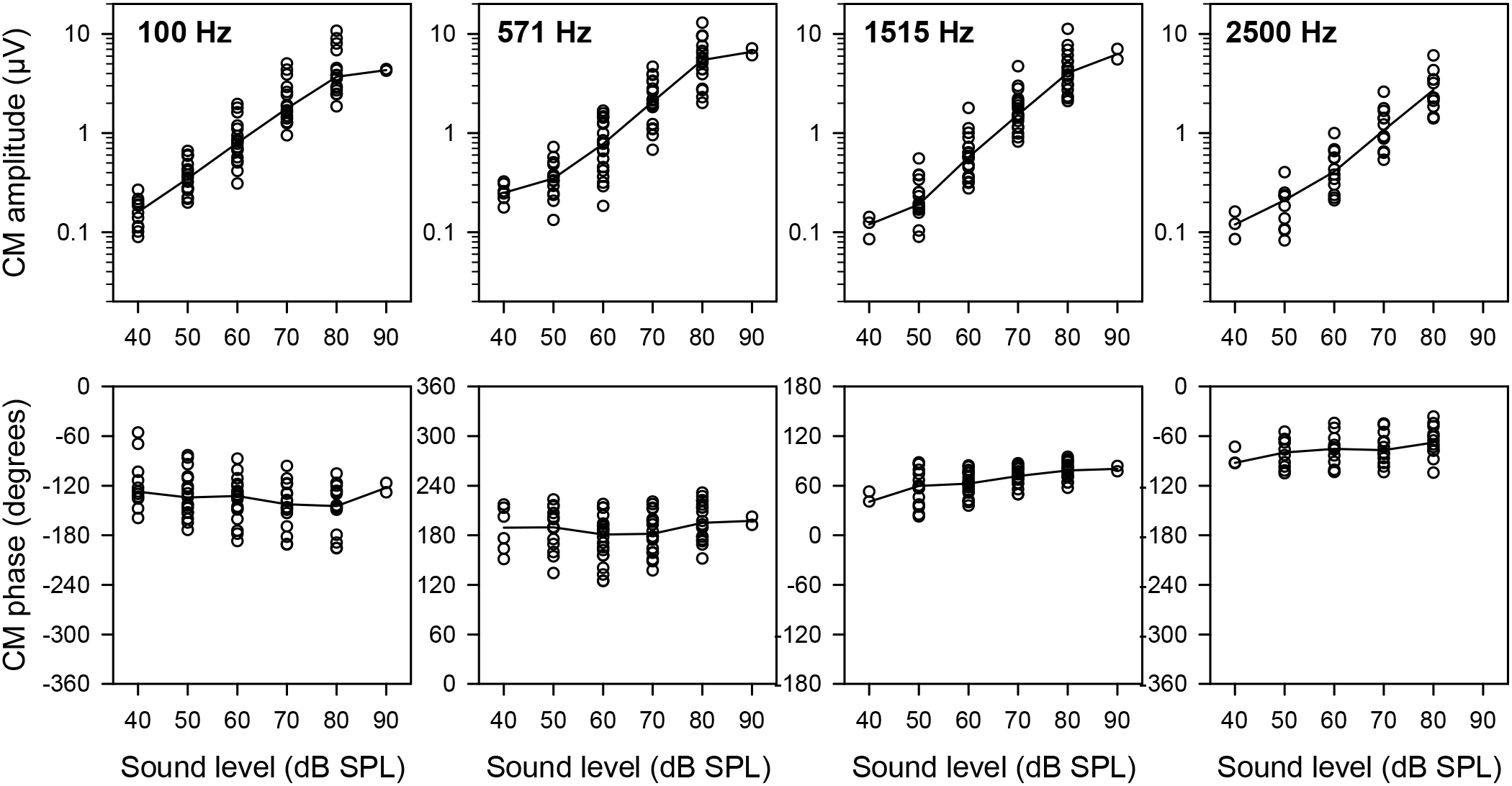
CM behavior with monaural ipsilateral stimulation of increasing sound level, at 4 different frequencies. Top row of panels: CM amplitude (in μV) as a function of level, at 100, 571, 1515 and 2500 Hz. Each panel shows raw data from both ears of 8 chickens, the solid line joins the median values at each level. Bottom row of panels: CM phase as a function of sound level.

**Fig. 2A:**
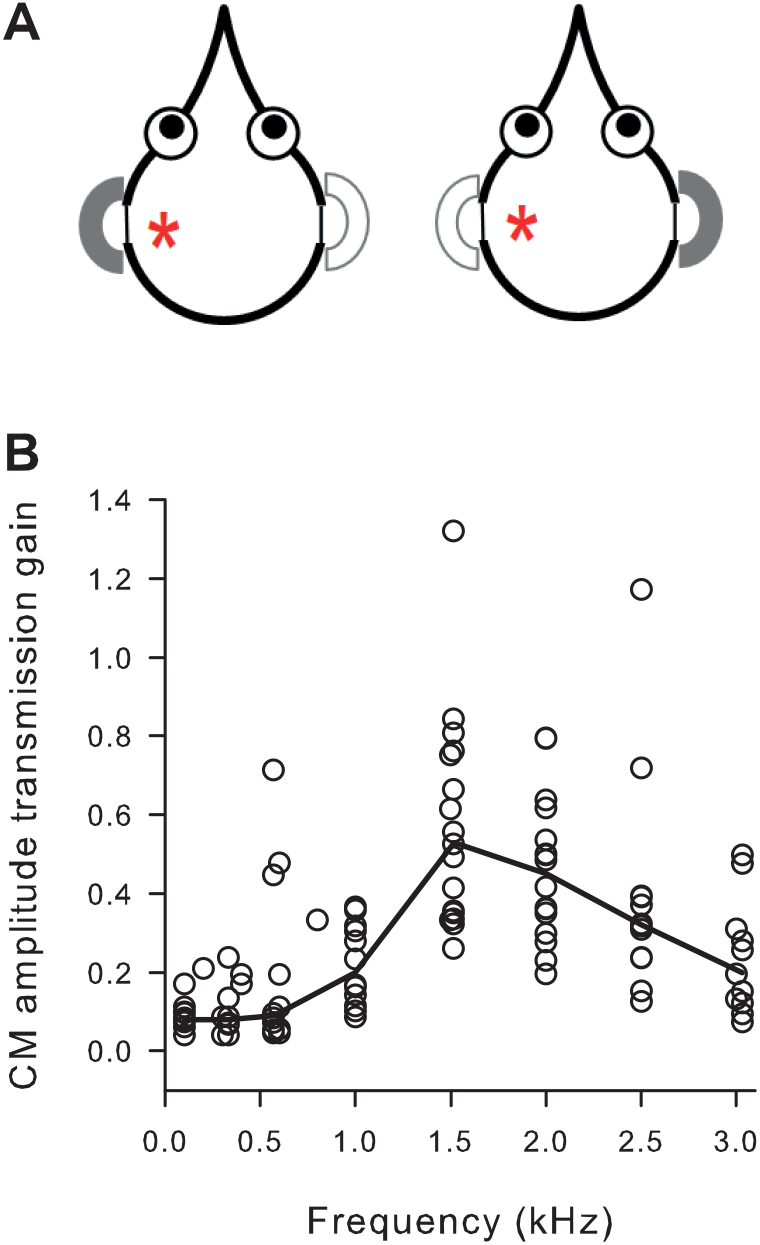
Cartoon illustration of same-ear comparison used to derive interaural transmission and delay. CM recordings from the same ear were compared (indicated by the red star), in response to monaural stimulation of the ipsi- or contralateral ear (indicated by solid gray earphone). B: Interaural transmission gain, i.e. the ratio of contra- to ipsilateral CM amplitude, at 70 dB SPL. Shown are individual measurements from both ears of 8 chickens. The solid line joins the median values at each standard frequency.

The phase of the CM was nearly invariant with level. Variations were not systematic and typically less than 30° over a 30 dB range. Examples are shown in Fig. 1, bottom row of panels.

### CM measurements of interaural transmission amplitude and delay

Interaural transmission was determined by comparing CM amplitudes from the same ear upon stimulation with 70 dB SPL from the ipsi- and contralateral side (Fig. 2A). Transmission was expressed as the ratio of contra- to ipsilateral CM amplitude, comparable to the amplitude transmission gain derived from eardrum vibration measurements (Michelsen and Larsen, 2008). Amplitude transmission gain was maximal, at a median value of 0.53, at 1.5 kHz (Fig. 2 B), corresponding to −5.5 dB attenuation. Minimal transmission, with gain values below 0.1 (equivalent to >20 dB attenuation), was observed at frequencies below 1 kHz.

Interaural delay was estimated in two different ways. First, a fixed delay should, with increasing frequency, result in a linearly rising phase accumulation in the contralateral CM. Indeed, the unwrapped plot of phase as a function of frequency was reasonably fit by a linear regression with a slope corresponding to a time delay of 264 μs (Fig. 3). The phase of the ipsilateral CM varied randomly over frequency, indicating no or only very small (acoustic and transduction) delays. Second, to examine more closely for any frequency dependence, the phase difference between the paired, same-ear CM measurements upon ipsi- and contralateral stimulation was determined and converted to the corresponding time delay. The phase of the contralateral CM was inverted by 180° before this comparison, to account for the fact that the same stimulus phase which causes inward motion of the ipsilateral eardrum will cause outward motion of the contralateral eardrum after travelling through the interaural connections, and will thus trigger an inverted CM response (Rosowski and Saunders, 1980; Larsen et al., 2006). With pure-tone stimulation, as used here, the phase comparison carries an inherent cyclic ambiguity. No assumptions were made about which side should be leading, thus the reported phase differences are minimal values. As expected, these phase differences showed a similar frequency dependence as the contralateral CM phase readings alone. However, after converting to time differences, the deviations from linearity became apparent as a systematic decrease of interaural delay with increasing frequency. There was an initial drastic decrease from nearly 4000 μs at 100 Hz to a median value of 380 μs at 1 kHz, and a subsequent shallower decline to a median of 264 μs at 3 kHz (Fig. 3B).

**Fig. 3:**
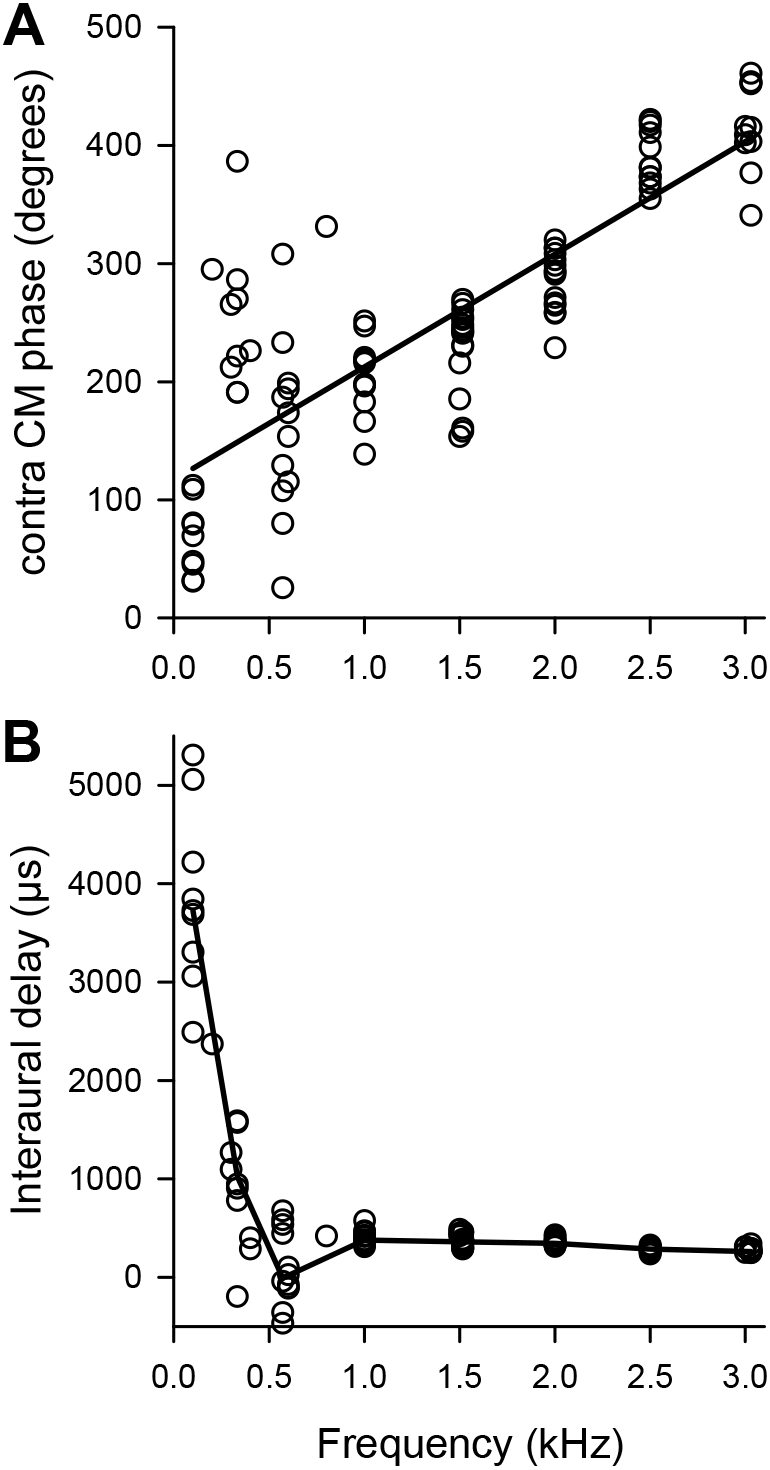
Measurements of interaural delay. A: Phase of the CM with contralateral stimulation at 70 dB SPL, unwrapped over different frequencies. Shown are raw data from both ears of 8 chickens. The solid line is a linear regression (y = 117.16 + 0.095x, r = 0.82, p<0.001, n = 99) the slope of which corresponds to a constant delay of 264 μs. B: Interaural delay derived from same-ear comparisons as illustrated in Fig. 2A. Phase differences were converted to time delays. The solid line joins median values at each standard frequency.

After blocking the interaural connections, the great majority of contralateral CM signals that had initially been above criterion disappeared into the noise (44 of 55, or 80%, over all frequencies and levels). The few that still met criterion showed both significant reductions in amplitude and significant phase shifts, compared to the unblocked condition (Wilcoxon tests, p < 0.01, n = 11). Ipsilateral CM amplitudes and phases were unaffected. After blockage, interaural attenuation was generally above 30 dB and independent of frequency. Careful dissection of the manipulated heads after the experiment showed that the injected petroleum jelly had accumulated behind the eardrum and from there primarily ventral. The connection that is commonly called the interaural canal (Larsen et al., 2016) had been filled to approximately the skull’s midline.

### Amplitude modulation of a given-ear CM with binaural stimulation of varying ITD

Upon binaural stimulation with equal sound levels, but varying ITD, CM amplitudes clearly modulated with ITD (example in Fig. 4A). This is the equivalent of the directionality of eardrum vibration shown with free-field stimulation (review in (Christensen-Dalsgaard, 2011). Without any internal coupling, both CM amplitudes are expected to remain unaffected by the varying ITD, as binaural sound levels were kept constant. If significant internal coupling exists, CM amplitudes are expected to modulate in an ITD-dependent fashion, as eardrum vibration would modulate as a function of azimuthal sound-source position in the free field. The extent of this modulation was quantified as the ratio of maximal to minimal amplitude over a range of ± one period of ITD at the respective test frequency, and termed the ITD modulation ratio. This ITD modulation ratio was equal in both ears (Wilcoxon test, p = 0.372, n = 115). However, it clearly varied with frequency. Maximal ITD modulation ratios occurred at 1.5 and 2 kHz, with median values of 2.22 and 1.99 (max. 6.01). Ratios decreased towards both lower and higher frequencies (Fig. 4B), mirroring the frequency dependence of interaural transmission. Median modulation ratios were consistently higher at 50 dB SPL as compared to 70 dB SPL. However, this difference was only significant for low frequencies. At 1 kHz and above, modulation ratios did not differ significantly with sound level (Fig. 4B; Mann-Whitney U-tests, p below or above 0.05, respectively).

**Fig. 4:**
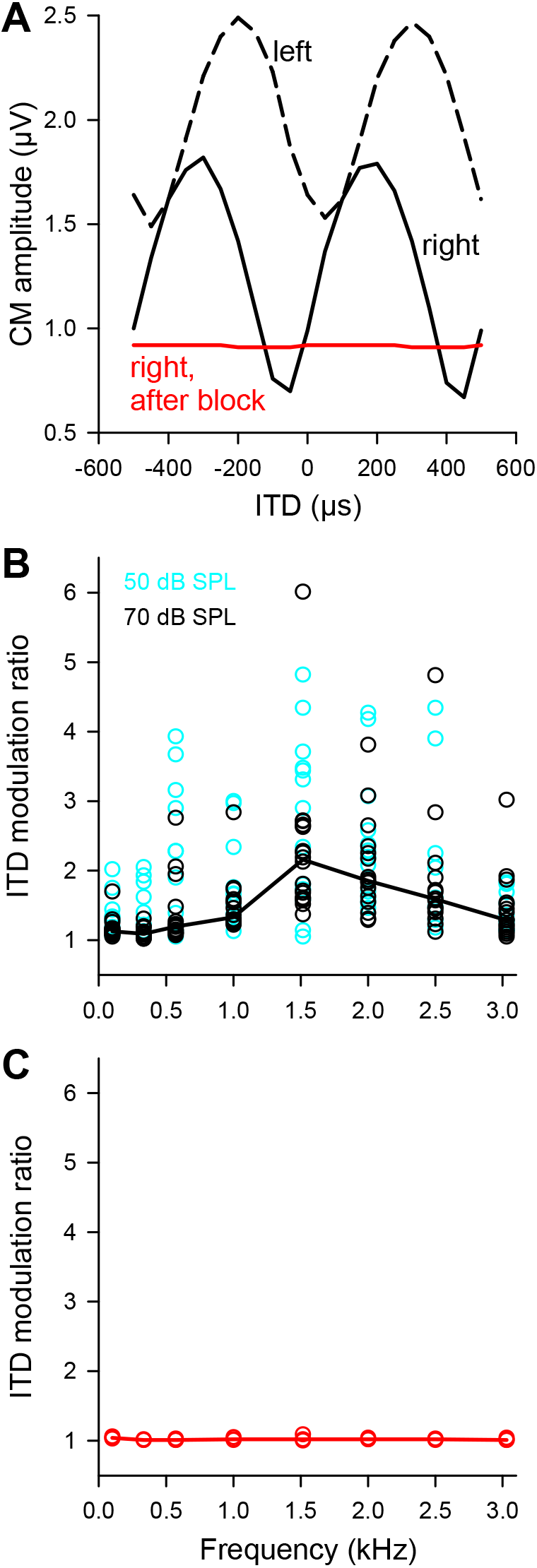
Modulation of CM amplitude upon binaural stimulation with varying ITD. A: Example of simultaneous CM recordings in both ears of an individual chicken, stimulated at 2 kHz and 70 dB SPL. Both CM recordings clearly modulated in amplitude as a function of ITD; modulation ratios were 1.63 (left) and 2.35 (right). Note the complete absence of modulation after blockage of the interaural connections (data shown in red; modulation ratio 1.02). B: ITD modulation ratios for all measurements in all ears (8 chickens), at two different sound levels, 50 dB SPL (blue circles) and 70 dB SPL (black circles). The solid line joins the median values for 70 dB SPL at each standard frequency. C: ITD modulation ratios at 70 dB SPL, after blockage of the interaural connections in 3 chickens.

Importantly, the modulation of CM amplitude with ITD was consistently abolished upon blockage of the interaural connections (Fig. 4A, C). Because our method of blockage from one side also impaired the ear ipsilateral to the manipulation, only the remaining good, contralateral ear could be evaluated. ITD modulation ratios in the remaining good ear never exceeded 1.09 at all frequencies (median values 1.01 – 1.04, Fig. 4C).

On average, the CM showed consistently higher maximal amplitudes and lower minimal amplitudes in the binaural condition, compared to monaural stimulation at the same sound levels. This suggests both constructive and destructive phase interference with binaural input. However, a frequency dependence was also obvious. CM maximal amplitudes at frequencies between 1 and 2.5 kHz to binaural stimulation at 70 dB SPL were reliably reduced after blockage of the interaural connections (same individual ears compared; only unmanipulated side; example in Fig. 4A). In contrast, the amplitude change was more variable for lower frequencies and at 3030 Hz, with 2 out of 3 ears actually showing enhanced amplitudes after blockage of the interaural connections, suggesting a predominantly destructive interaction in the normal binaural condition at those frequencies.

### Comparison of ITD presented to ITD heard

Next, we used the phases of simultaneously recorded left and right CMs under binaural stimulation to derive the actual ITD that the animal experienced, the “ITD heard”. This is the analogous comparison to that performed by neurons in the binaural nucleus laminaris (e.g., Ashida and Carr, 2011). Phase differences were disambiguated and unwrapped, assuming that the difference that corresponded most closely to the acoustically presented ITD was the correct one (examples in Fig. 5A, E). In other words, we assumed that the actual phase difference could not differ from the presented one by more than 180°.

**Fig. 5:**
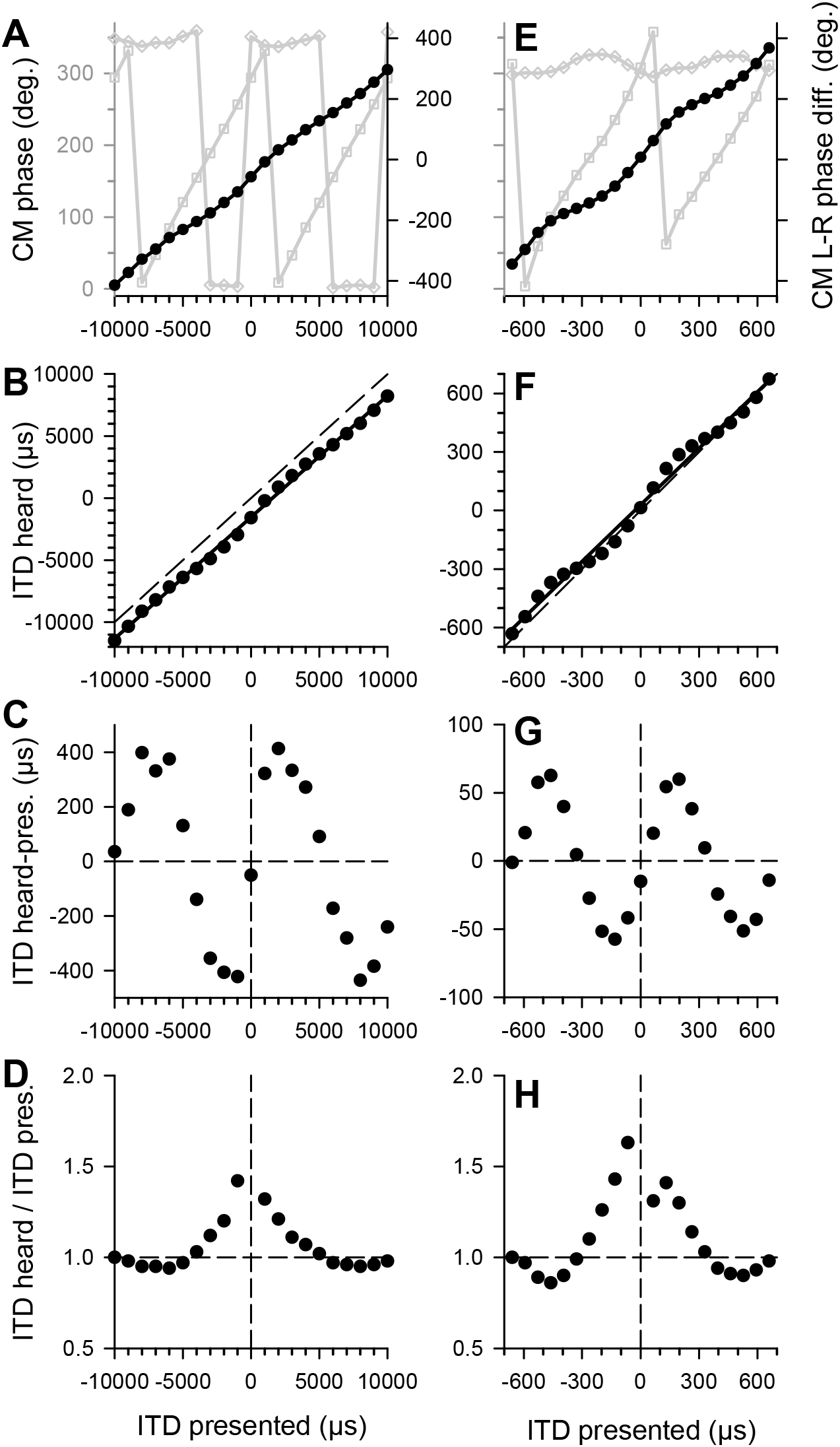
Two examples of the derivation of “ITD heard”. A-D: An example with binaural stimulation at 100 Hz, 70 dB SPL. E-H: An example with binaural stimulation at 1515 Hz, 50 dB SPL. The top panels (A, E) show the raw phases of the simultaneously recorded left and right CMs (grey symbols and lines, refer to left ordinate), as a function of the acoustically presented ITD, varied over ± one period of the stimulation frequency. Black symbols and lines represent the difference between the unwrapped left and right CM phases (refer to right ordinate). The next panels (B, F) show the phase differences converted to time difference, termed the ITD heard, as a function of the acoustically presented ITD. The solid lines are linear regressions to the data points, the dashed line indicates identical values for presented and heard ITD, for reference. Note that the data show both a constant offset and an ITD-varying deviation from this reference. The constant offset is represented by the y-axis intercept of the linear regression. For the data shown in the panels C and G, the constant offset has been subtracted, and the remaining deviation of the ITD heard from the ITD acoustically presented is shown as a function of the acoustically presented ITD. Note that the largest deviations occurred near the acoustic midline. Finally, panels D and H plot the ratio of ITD heard / ITD presented acoustically, for the same data.

The ITD heard commonly deviated systematically from the acoustically presented ITD. Many recordings showed two components to this: a constant offset from the expected (acoustically presented) ITD and an ITD-dependent deviation cycling at the period of the stimulation frequency. The constant offset was quantified as the y-axis intercept of the linear regression of ITD heard as a function of ITD presented (examples in Fig. 5B, F). The offset appeared to vary randomly within mostly ± 50 degrees (0.15 cycles), independent of frequency or sound level. However, there was a tendency for this offset to show a consistent polarity in a given animal. We therefore assumed it to be an artefact of slightly asymmetric recording conditions between the two ears. The offset was subtracted from all measurements and the unbiased difference between the ITD heard and the ITD presented was derived (examples in Fig. 5C, G). To highlight whether this deviation would have enhanced or reduced the perceived ITD relative to the acoustically presented ITD, the ratio between them was also determined (Fig. 5D, H). Note that ratios above 1 indicate a larger ITD heard, ratios below 1 a smaller ITD heard.

For both examples shown in Fig. 5, the largest ratios occurred around zero ITD, suggesting that the deviations would act to enhance ITDs in the chicken’s natural range. This was also typical at the population level. Figure 6 shows median data for 4 frequencies, at both sound levels tested, 50 and 70 dB SPL. Median ratios were generally positive around the acoustic midline, with the exception of 333 Hz (Fig. 6, second row), where the ratios were negative, suggesting an unfavourable compression of the ITD range heard. A further, unexpected observation was that the extent of enhancement (or compression, at 333 Hz) could be level-dependent. Ratios were often, but not universally, higher at 50 dB SPL than at 70 dB SPL (Fig. 6). The highest median ratio, 1.86 at 1515 Hz and 70 dB SPL, suggested an enlargement of the ITD heard by a factor of 1.8, compared to the acoustically presented ITD.

**Fig. 6:**
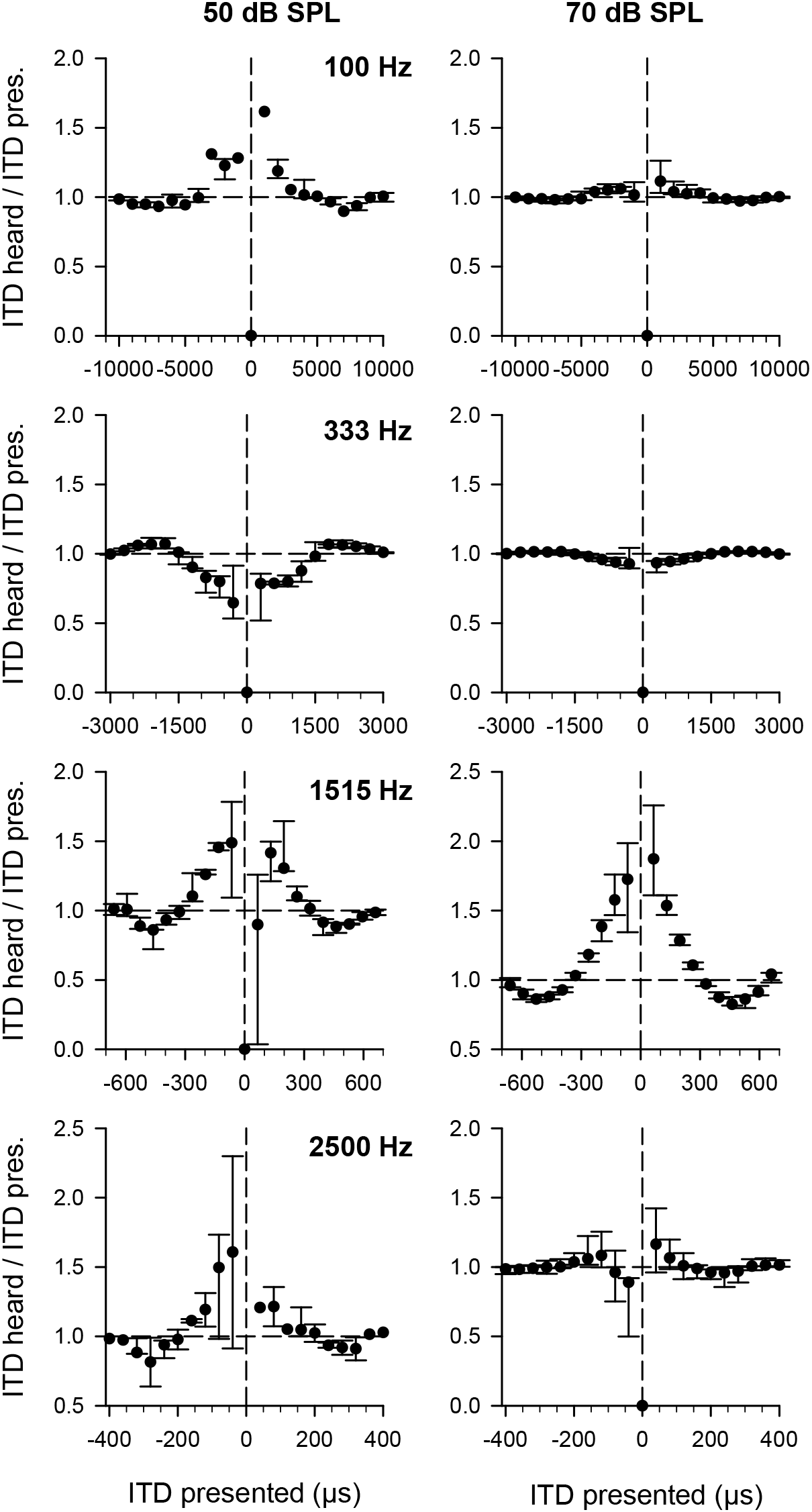
Median ratios of ITD heard / ITD presented acoustically. Data for 4 different frequencies are shown: 100, 333, 1515 and 2500 Hz, in successive panel rows. The two columns of panels show data for two different sound levels: 50 dB SPL (left) and 70 dB SPL (right). Medians and interquartile ranges are plotted as a function of the acoustically presented ITD. Vertical dashed lines indicate the acoustic midline, i.e. zero ITD, and horizontal dashed lines indicate identical values for ITD heard and ITD presented, i.e. a ratio of 1, for reference. Note that ratios above 1 indicate a larger ITD heard, ratios below 1 a smaller ITD heard. Note that this kind of plot highlights whether a deviation would increase the perceived ITD range (ratios > 1) or compress it (ratios < 1). At 100, 1515 and 2500 Hz, the effect was an enhancing one. However, at 333 Hz, the effect was compressive. Note also that at most frequencies, ratios were greater at the lower sound level of 50 dB SPL.

Finally, a prediction was derived from these data about the ITDs that the chicken should hear when a sound source originates in the free field from 90° to one side. For this, a value for the maximal acoustic ITD between the chicken’s ear canals needed to be chosen. According to the spherical head model of (Kuhn, 1977), an acoustic ITD of 100 μs should occur for chickens with a head width of 23 mm (as used here), or 130 μs for adult chickens with 30 mm head width. Acoustic ITD actually measured were around 170 μs for adult chickens (Schnyder et al., 2014; estimated from phase measurements shown in their Supplemental Fig. 9). As a best educated guess, we then calculated ITDs heard for 130 μs acoustically presented ITD. The prediction was derived by linear interpolation between adjacent data points, averaging ispi- and contralateral leading ITDs (i.e., assuming symmetry), and finally averaging the predictions derived from measurements at 50 and 70 dB SPL. Figure 7 shows the result together with previously published data (see Discussion).

**Fig. 7:**
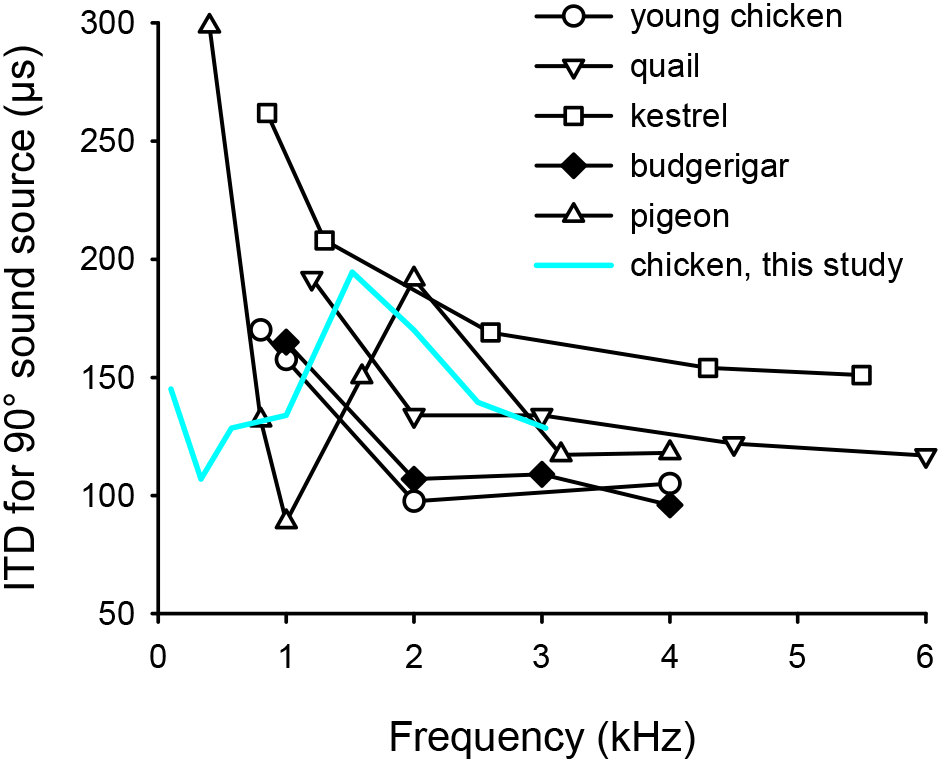
ITD heard from a sound source 90° to one side of an animal in the free field, as a function of frequency. Data shown in black are from published sources, distinguished by different symbols: Quail (head width 24 mm; Calford and Piddington, 1988), nankeen kestrel (31 mm; Calford and Piddington, 1988), young chicken (17 mm; Hyson et al., 1994), pigeon (22 mm according to Lewald, 1990; data shown are from Rosowski, 1979), and budgerigar (16 mm; Larsen et al., 2006). In blue, is shown a prediction from the present data, obtained with closed-system stimulation, by assuming a uniform acoustic ITD of 130 μs between the two ear canals. Any deviation from a flat line in such a plot suggests significant internal coupling of middle ears. Note the very similar trends of all datasets at frequencies above 1.5 kHz, but the much larger variation at lower frequencies, most prominently the markedly reduced ITD enhancement with closed-system stimulation in the present study.

**Fig. 8:**
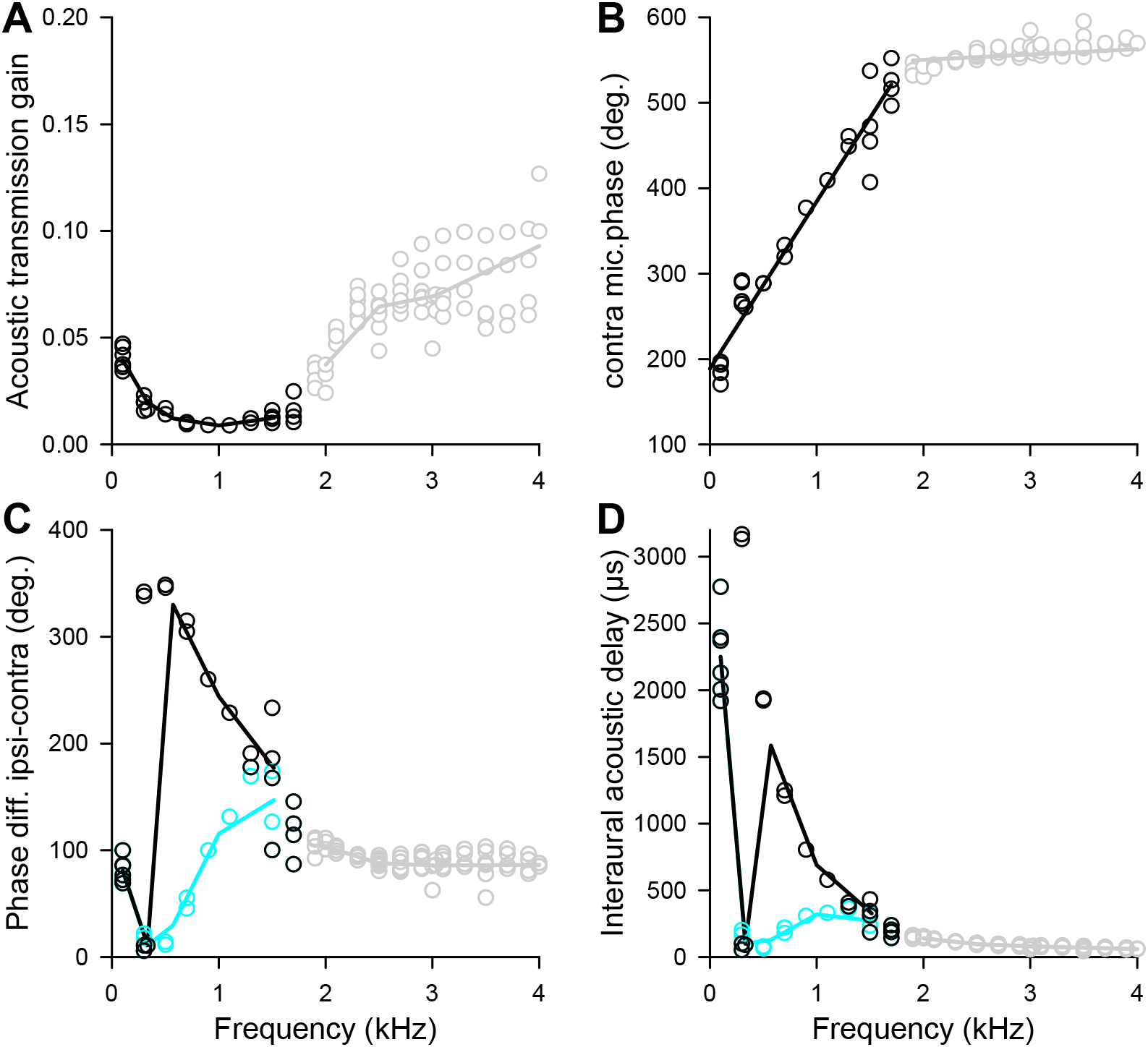
Measurements of acoustic interaural transmission and delay, using microphones in the outer ear canals, in a subset of 3 chickens. A: Acoustic transmission gain, derived in an analogous way to the CM data shown in Fig. 2B. B: Phase accumulation at the microphone contralateral to the simulation, analogous to the CM data shown in Fig. 3A. Note that above 1.8 kHz, virtually no phase accumulation occurred, suggesting direct electrical cross-talk between microphone channels. This data range is therefore shown grey in all panels. The solid lines represent linear regressions to the data below and above 1.8 kHz, respectively. The slope of the low-frequency regression corresponds to a constant delay of 544 μs. C: Phase difference between the microphone readings with monaural stimulation from the ipsi- or contralateral ear. Two different analyses are shown: either taking the minimal phase difference (blue symbols and line) or assuming that there should be a consistent ipsi lead and phase roll-off across frequencies (black symbols and line). The solid lines join median values at each standard frequency. D: The phase differences from C, converted to interaural time delays.

### Acoustical measurements of interaural transmission amplitude and delay

Acoustic measurements were derived in three chickens, using the microphones integral to the closed sound systems. These microphones were coupled to calibrated probe tubes that opened at the entrance to the chicken’s ear canal. Analogous to the CM analysis, interaural transmission was determined by comparing the readings from the same microphone upon stimulation with 70 dB SPL from the ipsi- and contralateral side, respectively. Transmission was again expressed as the ratio of contra- to ipsilateral amplitude reading. Acoustic interaural transmission was very consistent across animals but much lower than that shown by the CM measurements. Median values remained below 0.1 (equivalent to >20 dB attenuation) at all frequencies. Acoustic measurements also showed a different frequency dependence, with minimal transmission between 571 and 1515 Hz, and slightly rising towards both lower and higher frequencies (Fig. 7A). Measurements above 1.7 kHz were likely contaminated by artefacts (see next paragraph) and are thus shown in grey.

Interaural acoustic delay was first estimated from the slope of the phase accumulation across frequency at the contralateral microphone. This very clearly showed two components: a linear phase accumulation corresponding to a delay of 544 μs at frequencies up to 1.7 kHz, followed by a break to a much shallower slope, corresponding to a delay of only 17 μs at frequencies above 1.7 kHz (Fig. 7B). This suggests direct electrical pick-up across channels at higher frequencies. The value at lower frequencies was likely a truly acoustic delay.

Secondly, interaural acoustic delay was determined from the phase difference between both microphone readings. Only frequencies up to 1515 Hz were included, to minimize the influence of electrical cross-talk shown above. We tried to resolve cyclic ambiguity by measuring down to a very low frequency of 100 Hz, where the period was expected to far exceed the interaural delay. Furthermore, a fixed interaural delay should result in a linearly rising phase difference with increasing frequency. Lastly, it was assumed that the phase reading upon ipsilateral stimulation should lead that with contralateral stimulation.

However, the data did not clearly conform to those expectations. At 100 Hz, the ipsilateral readings consistently led the contralateral ones by a median of 81 degrees, corresponding to a delay of 2250 μs (Fig. 7C, D). However, assuming a continuing ipsilateral lead yielded phase differences which smoothly decreased (instead of increased) towards higher frequencies (Fig. 7C, black data). This translated into a corresponding decrease in interaural acoustic delay, down to a median value of 327 μs at 1515 Hz (Fig. 7D, black data). Abandoning the assumption of an ipsilateral lead and taking the shorter of the two possible leads in each case, led to a highly nonlinear phase-frequency relation (Fig. 7C, blue data). This translated to generally shorter interaural acoustic delays, between 100 and 330 μs, except at 100 Hz, where the median remained at 2250 μs (Fig. 7D, blue data).

After blocking interaural connections, at frequencies up to 1.7 kHz, contralateral microphone readings that had shown a significant signal in the unblocked condition dropped below criterion in nearly half the cases (7 of 18). Readings that remained above criterion did not show a mean change in either level or phase (Wilcoxon test, P > 0.05, n = 11). At higher frequencies, all contralateral microphone readings remained essentially unchanged after blocking interaural connections, which is consistent with the above conclusion of electrical pick-up. Even disregarding the higher frequencies, these observations nevertheless suggest that the blockage of interaural connections was either incomplete or that significant other sound paths existed.

## Discussion

The present study obtained clear evidence for a significant modulation of the sound localization cue ITD experienced by the chicken, relative to that presented acoustically to each ear. This modulation was shown to be mediated by the physical coupling between the middle ears, because blocking the interaural connection abolished the modulation. Data from zebra finch, pigeon and alligator suggest the presence of several distinct connective pathways across the head, including the most easily identified, ventrally directed interaural canal (Bierman et al., 2014; Larsen et al., 2016; Rosowski, 1979). The blocking experiments reported here are consistent with the existence of additional pathways in the chicken, too. Visual inspection suggested that the block typically filled the space immediately behind the eardrum and the ventral interaural canal, on the injected side. Although this largely eliminated all ipsilateral CM responses and bilateral CM recordings were no longer possible after the block, the acoustic measurements by microphones in the ear canals indicated some remaining crosstalk.

Some previous studies had already suggested that ITD was being significantly modulated by internally coupled middle ears, both in chickens (Hyson et al., 1994) and other avian species (Calford and Piddington, 1988; Larsen et al., 2006; Rosowski, 1979). Other studies, however, remained unconvinced of any significant physiological coupling (Klump and Larsen, 1992; Lewald, 1990). The specific value added by the present study is severalfold: 1) The chicken is a popular model species in auditory localization research. These are the first measurements of interaural transmission and delay in chickens that consciously avoided the confounding artefact of negative pressure buildup in the middle ear under anesthesia. 2) By using cochlear microphonics, most of the frequency range that is relevant to the chicken and many other birds could be probed, including low frequencies down to 100 Hz. 3) ITD was determined in the same individuals. By stimulating through closed sound systems, a situation typically used in neurophysiological tests for ITD selectivity was replicated.

### Validity of CM measurements as a proxy for eardrum vibration

Different methods have been employed to experimentally verify the effect of internally coupled ears. Arguably the most elegant and direct way is to measure eardrum vibration in the intact animal, using laser Doppler vibrometry (review in Michelsen and Larsen, 2008) which, ideally, avoids any kind of invasive manipulation. However, an important limitation is the often inadequate signal-to-noise ratio at low frequencies, below 1 to 2 kHz. This excludes a substantial part of the frequency range of interest in the debate about ITD cues and their neural coding. Furthermore, aiming a laser beam onto the eardrum is, in practice, difficult to combine with the use of closed sound systems. Measurements of CM, on the other hand, while not suffering the above restrictions, are only an indirect correlate of eardrum motion. Although it is undisputed that hair-cell responses are the principal source of the CM, the source distribution within the cochlea upon stimulation with different frequencies is not well characterized in birds (Köppl and Gleich, 2007).

An important prerequisite to using the CM as a proxy for eardrum vibration is that it behaves linearly within the SPL range of measurements. This was satisfied here. CM amplitudes grew linearly with sound level up to 80 dB SPL (Fig. 1, top row of panels). Important for phase comparisons, the phase of the CM was, on average, invariant with level (Fig. 1, bottom row of panels), consistent with the findings of Calford and Piddington (1988) in quails. Small nonlinearities are difficult to exclude and are the likely cause for the minor level dependencies observed in binaural data (Fig. 6). One likely source of nonlinearity is a different (larger) set of hair-cell generators at higher sound levels. Another possibility is efferent feedback to the hair cells which could conceivably occur within the analysis window used here (Kaiser and Manley, 1994). In contrast, the middle-ear reflex is only triggered during vocalization in chickens (Counter and Borg, 1979; Larsen et al., 1997) and was thus not likely in the present experiments.

### Comparison with previous estimates of interaural transmission and delay in birds

Previous studies in different bird species did not universally agree on the principle existence of significant internal coupling between the middle-ear spaces. A large part of the variation between studies is likely due to two experimental artefacts that reduce interaural transmission in a frequency-specific manner, as compared to the natural situation of an awake bird in the acoustic free field. One of these detrimental conditions is the potential build-up of negative middle-ear pressure in anaesthetized birds (Larsen et al., 2016; Larsen et al., 1997). The occurrence and extent of this artefact are highly variable and species-specific and may thus have led to decreased estimates of interaural coupling in earlier studies (lack of awareness of the problem) to unknown degrees. Indeed, interaural transmission values obtained in awake birds or under anesthesia but with middle-ear ventilation ensured, tend to be the highest reported: around a maximal gain of 0.55 or −5 dB attenuation (Larsen et al., 1997) and 0.3 or −10 dB (Larsen et al., 2006) for anesthetized and awake budgerigars, respectively, 0.5 or −6 dB in the anaesthetized barn owl (Kettler et al., 2016), and 0.53 or −5.5 dB in the present study for anesthetized chickens.

Furthermore, there is evidence that sealing closed sound delivery systems to the ear canal(s) also acts to reduce interaural transmission, and also disproportionately at lower frequencies. Although this is difficult to disentangle from the middle-ear pressure artefact in older work, the present study adds considerable strength to that hypothesis. Using closed sound systems sealed to both ear canals, two previous studies, in chicken (Rosowski and Saunders, 1980) and pigeon (Rosowski, 1979), as well as the present study observed relatively less interaural transmission at low frequencies. In the starling, interaural transmission was flat up to 3.5 kHz, with only one ear canal sealed to a closed sound delivery system (Klump and Larsen, 1992). In contrast, with open-field stimulation, interaural transmission in the budgerigar was most effective at about 1 kHz, compared to frequencies above that (Larsen et al., 2006; Larsen et al., 1997). In dead quail (where the above anesthesia artefact should not have occurred), a direct comparison of free-field and closed-field stimulation showed the same relative reduction of transmission at low frequencies, with one ear canal sealed to a closed sound delivery system (Hill et al., 1980). Such changes under headphone conditions may be due to restricting the air volume coupled to the external auditory meatus and thus changing middle-ear stiffness, similar to what has been shown in frogs (Gridi-Papp et al., 2008; Pinder and Palmer, 1983).

Measurements of interaural delay across the head, i.e. the transmission time for sound between an ipsilateral source and the inside of the contralateral eardrum, typically show values that are clearly larger than the acoustic travel time across the linear head width. In chickens, budgerigars, starlings and barn owls, phase measurements of eardrum vibration or CM yielded estimated interaural delays of 70 to 232 μs, which correspond to 2 to 4 times the equivalent interaural distances of those birds (Kettler et al., 2016; Larsen et al., 2006; Rosowski and Saunders, 1980). The present mean value of 264 μs interaural delay for the chicken also falls within this range, and corresponds to nearly 4 times the equivalent head width of the chickens used. Perhaps most strikingly, the interaural delay in the chicken was frequency dependent, with values increasing into the millisecond range at the lowest frequencies evaluated here, and similar for both acoustic and CM measurements. Two previous studies, also using closed-system stimulation, extended to similarly low frequencies. In pigeon CM and acoustic measurements, Rosowski (1979) found a very similar frequency dependence (converting his phase values to time), with maximal delays of about 600 μs at 160 – 200 Hz, and around 120 μs above 1 kHz. Acoustic measurements in chickens, however, with the identical technique, found no interaural delay at all for frequencies up to 1 kHz (Rosowski and Saunders, 1980). Clearly, these data sets cannot be reconciled and currently remain unexplained. Evidence that multiple sound paths across the avian skull exist, have recently led to speculations about how these different paths might interact and create frequency-dependent phase shifts (Larsen et al., 2016).

In summary, interaural transmission and interaural delay are salient parameters that determine what exactly arrives at the contralateral eardrum after traversing the head. All the available data agree that sound does not simply travel unimpeded across the avian head but is attenuated and significantly delayed. The degree of interaural transmission has been underestimated so far in birds and probably typically peaks for low frequencies around a gain of 0.5, or −6 dB attenuation. Although this gain is not as high as in lizards who hold the record of nearly unimpeded interaural transmission (Christensen-Dalsgaard and Manley, 2008), it is of the same order as in frogs and insects (Christensen-Dalsgaard, 2011; Michelsen and Larsen, 2008) and should put to rest any remaining doubts about the significance of internal coupling between avian middle ears. However, it is important to emphasize that under closed-field stimulation, as used here and typically in neurophysiological experiments, interaural transmission at low frequencies is likely compromised. The interaural delay is typically several times longer than expected from simply traversing the head width, consistent with anatomical evidence for complex sound paths through the avian skull. Any frequency dependence of the interaural delay and possible artefactual alterations remain ill-characterized and this still makes it in particular difficult to predict the ITD resulting from internal coupling of avian middle ears.

### Extent of binaural ITD enhancement

The present data showed significant internal coupling between the chicken’s middle ears. Furthermore, our measurements clearly suggested an expansion of the ITD range heard, compared to what was acoustically presented with binaural stimulation, at some of the frequencies evaluated. However, data obtained under closed-system headphone stimulation do not directly translate to free-field conditions. As discussed above (see previous section), interaural transmission is likely compromised with closed-field stimulation at low frequencies, thus underestimating the potential enhancing effects on ITD. On the other hand, when changing the position of a sound source under free-field conditions, ILDs occur in addition to ITDs, while in our headphone experiments, ITDs were presented in isolation. This will tend to maximize interaural effects, since the sound of a simulated contralateral source is then only attenuated by the interaural connections and not, in addition, by head and body shadowing. However, compared to the interaural attenuation, the attenuation by diffraction is the minor component at frequencies up to about 4 kHz (Larsen et al., 2006).

Figure 7 validates those assumptions. Here, the prediction from our data, of ITD heard from sound sources originating 90° to one side, is shown together with published ITDs derived from CM or eardrum vibration recordings under free-field conditions, for sound sources 90° to one side, in birds with approximately similar head sizes: Quail (head width 24 mm; Calford and Piddington, 1988), nankeen kestrel (31 mm; Calford and Piddington, 1988), young chickens (17 mm; Hyson et al., 1994), budgerigar (16 mm; Larsen et al., 2006), and pigeon (22 mm; Rosowski, 1979). As expected, the data obtained under free-field conditions mostly show larger ITDs at low frequencies, below 1 kHz. At higher frequencies, however, our data are a good match. The comparison supports the notion that, 1) under natural free-field conditions, ITDs are enhanced by the internally coupled middle ears, 2) increase with decreasing frequency, and 3) reach at least 200 μs in a bird of adult quail or chicken size. The low-frequency range, below 1 kHz, still shows the largest uncertainties. Currently, it can only be assumed that the ITD heard continues to rise with decreasing frequency, but the precise value of the increase remains unknown. Eardrum vibration data do not extend to such low frequencies, since the velocity measurements typically used are insufficiently sensitive. In addition, the well-defined free-field presentation of such low frequencies requires large anechoic chambers, which may explain why most of the classic CM measurements using free-field stimulation also did not probe such low frequencies. The present study demonstrated that the use of headphone stimulation is also not an alternative, because this in itself alters the properties of the internal coupling.

### Implications for the interpretation of neural recordings

One main motivation for the present study was to clarify the influence of the internally coupled middle ears of chickens under the standard experimental conditions used during neurophysiological recordings from neurons involved in ITD processing. In such experiments, acoustic stimulation through closed sound systems sealed to both ear canals is the norm because 1), it enables controlled, separate stimulation of the two ears, allowing, e.g., to vary only ITD, in order to probe the specific selectivity of neurons and 2), a well-defined acoustic free field is difficult to achieve due to the extensive equipment necessary for invasive neurophysiology and typically surrounding the experimental animal (Michelsen and Larsen, 2008).

An important lesson from the present study for the interpretation of neurophysiological data is that the ITD that is acoustically played by the headphones is not necessarily what is relayed by the two inner ears and subsequently compared by the binaural brainstem neurons. In other words, neurophysiological responses are referred to the wrong ITD in such cases. Furthermore, for tonotopically organized nuclei in which the individual neurons are also narrowly frequency tuned, the errors introduced may differ between frequency ranges. The present data suggest that, in the chicken, the ITD range responded to by neurons with best frequencies between approximately 1.5 and 2.5 kHz will be artificially compressed, because the ITD heard is significantly larger than that acoustically presented. Conversely, responses of neurons around 300 Hz will show artificially inflated ITD ranges, because here, the ITD heard under headphone conditions is actually smaller than that acoustically presented.

One might argue that the errors introduced in that way are small and should not affect principal findings. However, the debate about what constitutes a physiologically meaningful ITD response in binaural neurons has a particular and controversial history (e.g., Joris and Yin, 2007; McAlpine, 2005). Some of it was based on incorrect (too low) assumptions about the naturally heard ITD-range of animals, both for mammals and birds (see Introduction). The present study has identified an additional confounding factor in animals with internally coupled middle ears, i.e. non-mammalian species. Unfortunately, the present results cannot be assumed to generalize quantitatively to other species, i.e. the specific artefacts introduced by headphone stimulation need to be identified in each case.

## Acknowledgements

I thank Mark Konishi for the generous gift of the software “xdphys” custom-written in his lab and used in these experiments, and STELS-OL (Scientific and Technical English Language Services, Oldenburg, Germany, stels-ol.de) for English language editing.

## No competing interests declared

